# Expansion of Olfactory Receptor Family 14 Across Fossorial and Insectivorous Mammals

**DOI:** 10.64898/2026.07.15.738794

**Authors:** Louise Ryan, Graham M. Hughes

**Affiliations:** School of Biology and Environmental Science, University College Dublin, Ireland

**Keywords:** OR14, Olfactory Receptors, Gene Family Expansion, Mammalian Evolution, Insectivory, Fossorial, Sensory Perception

## Abstract

Mammals have incredibly diverse olfactory repertoires, reflected in significant copy number variation of olfactory receptor genes across species. As mammals occupy a wide range of habitats and foraging environments, it is expected that such variation is shaped by ecological niche. Indeed, several studies have shown that specific olfactory receptor gene families are associated with habitat and dietary specialization. Despite this, there is currently little known about the ecological factors driving variation of olfactory receptor family 14 (OR14) across mammals.

Here, we address this gap by mining mammalian genomes to identify and classify OR14 genes across hundreds of species. By mapping OR14 variation onto dietary and habitat variables, in the context of the mammalian phylogenetic tree, we identify key factors associated with OR14 evolution across mammals.

Specifically, we find that OR14 has undergone repeated expansion across independent lineages of fossorial and insectivorous mammals, suggesting that expansion of this gene family may be adaptive and driven by convergent ecological pressures. By performing principal component analysis and k-means clustering based on OR14 subfamily composition, we find that species cluster irrespective of phylogeny. Furthermore, we show that many independent myrmecophagous lineages form a single cluster, providing evidence of convergence at the sequence level, potentially driven by selection for ant/termite rich diets. Together, our results indicate that OR14 plays an important functional role in fossorial and insectivorous mammals, providing a foundation for future studies aimed at deorphaning these receptors.

## Introduction

Mammals are a diverse group of vertebrate species that exhibit vast morphological adaptations and occupy almost every ecological niche on the planet (Foley et al. 2016). Characterized by the presence of mammary glands, fur, and endothermic systems, mammals have occupied several habitats (e.g. terrestrial, aquatic, volant, fossorial), and dietary niches (e.g. carnivory, frugivory, insectivory) throughout their evolutionary history (Foley et al. 2016, Kissling et al. 2014, Solari and Baker 2007, Wilman et al. 2014). A key feature of mammalian evolution is the modification of sensory perception, as the ability to detect novel odors, sounds, and tastants is crucial for adaptation to novel environments. Consequently, convergent signals of sensory adaptation are evident throughout mammalian evolution. For example, echolocation has independently evolved in bats and odontocetes, allowing them to navigate in complete darkness and in marine environments, respectively (Liu et al. 2010, Parker et al. 2013). Similarly, complete loss or degradation of visual systems is observed across independent fossorial lineages, which rely instead on tactile or chemosensory cues for navigation in low-light subterranean environments (Crish et al. 2003, Heth et al. 2002, Partha et al. 2017, Springer et al. 2023).

At the molecular level, sensory perception is mediated by sensory receptor gene families. These receptors exhibit significant copy number variation across species (Dong et al. 2009, Francia et al. 2015, Lagman et al. 2013, Nishihara et al. 2024, Policarpo et al. 2024, Ryan and Hughes 2026, Vallender et al. 2010, Young et al. 2009), often reflecting adaptation to ecological niche (Hayden et al. 2010, Hayden et al. 2014, Hughes et al. 2018, Jiang et al. 2012, Niimura 2009, Niimura and Nei 2005, Zhao et al. 2010a, Zhao et al. 2010b). This is particularly true of olfactory receptors (ORs), which form one of the largest multigene families in mammals (Niimura et al. 2014). Expressed in the olfactory epithelium of the nasal cavity, OR repertoires range from tens of functional genes in aquatic species, to hundreds in humans, and thousands in species such as mice and elephants (Hughes et al. 2018, Niimura et al. 2014, Policarpo et al. 2024, Ryan and Hughes 2026). Based on phylogenetic reconstruction, ORs can be further subdivided into two classes, class I and class II, each consisting of multiple OR families. Class I includes the OR51, OR52, OR55, and OR56 families, while OR1/3/7, OR2/13, OR4, OR5/8/9, OR6, OR10, OR11, OR12, and OR14 belong to class II (Hayden et al. 2010, Hughes et al. 2018). Each OR family contains multiple subfamilies, with over 600 subfamilies in total reported in the literature (Olender et al. 2020).

Investigating patterns of expansion and loss across OR families can provide insight into the ecological factors driving olfactory tuning. Several studies have linked lineage-specific changes in OR family size to habitat and dietary niche (Hayden et al. 2010, Hayden et al. 2014, Hughes et al. 2018). For example, Hayden et al. (2014) uncovered a link between frugivory and expansion of the OR1/3/7 family in bats. This observation is further supported by Hughes et al. (2018), which show that species-specific duplications within OR1/3/7 are significantly associated with frugivorous diets across 58 mammalian species. Despite growing evidence that environmental variables influence OR family evolution, the ecological factors driving variation across specific OR gene families, such as OR14, remain unknown. Coupled with the fact that very few ligands have been characterized for OR14 receptors (Ollitrault et al. 2024), the functional relevance of this specific OR gene family across mammals remains ambiguous.

Previous studies exploring the evolution of the OR14 gene family have highlighted unique expansion in monotremes and marsupials (Sampson et al. 2024) and a high number of functional receptors in pangolins (Li et al. 2024) and certain rodent lineages (Courcelle et al. 2023). However, there has yet to be a large-scale investigation into the expansion of this gene family in light of the ecological factors underlying variation across a broader taxonomic sample. Here, we classify the OR14 gene family across 555 mammalian species with the aim of characterizing lineage-specific expansion and loss across the mammal species tree. By coupling variation in OR14 repertoires with dietary and habitat ecological niches, we show that repeated OR14 expansions are associated with fossoriality and insectivory and explore whether divergence across OR14 subfamilies reflects ecological specialization.

## Methods

### Identification and Classification of OR14 Genes across Mammals

Reference genome assemblies across 555 mammalian species were downloaded from the NCBI RefSeq (O’Leary et al. 2016) and Genbank (Sayers et al. 2024) databases (Table S1). OR mining and gene prediction was performed using Sensommatic (Ryan et al. 2024). Default parameters were selected for each run, with ‘human’ chosen as the AUGUSTUS (Stanke et al. 2004) training species. Receptors less than 860 bp in length were considered truncated and classified as pseudogenes, as they were unlikely to encode a complete seven transmembrane domain (Ryan et al. 2024).

To further classify ORs at the family level, the Maximum Mutual Similarity (MMS) algorithm (Olender et al. 2020) was applied, using classifications from the HORDE database (Olender et al. 2013) to guide family assignment. Prior to classification, nucleotide sequences were translated to amino acid sequences. FASTY-36 (Pearson and Lipman 1988) was used to account for frameshift mutations when translating pseudogenes. Receptors sharing ≥40% amino acid sequence identity with a receptor on the HORDE database (Olender et al. 2013) were assigned to the corresponding OR family. Unclassified receptors (<40% identity) were discarded from downstream analyses. Non-chromosomal receptors were excluded from the human repertoire to remove allelic variants. The total number of functional and pseudogenized OR14 genes were quantified for each species (Table S1). To account for general OR expansion, the proportion of functional OR14 genes was also quantified, expressed as a percentage of the total number of functional ORs in each species. Based on manual inspection, one OR14 gene in *Eidolon helvum* (Straw-colored fruit bat) was misclassified as a pseudogene due to an erroneous truncated prediction. Manual sequence correction of this receptor was performed with TBLASTN (Altschul et al. 1990), using an ortholog from the sister taxon *Eidolon dupreanum* (Malagasy straw-colored fruit bat) as the query (Figure S1).

### Phylogenetic Mapping of OR14 Repertoires and Ecological Traits

To visualize copy number variation of OR14 genes across mammals, TimeTree5 (Kumar et al. 2022) was used to generate a mammalian species tree representing 529 of the 555 species analyzed. Using the *ggtree* R package (Yu 2020), functional OR14 gene counts and relative OR14 proportions were mapped onto the phylogenetic tree (Figure 1). To explore which factors influence OR14 evolution, dietary and ecological trait data were compiled from public repositories and mapped to the tree (Table S2). The percentage of invertebrates consumed as part of each diet were obtained from the EltonTraits database (Wilman et al. 2014). Foraging strata, detailing where in a habitat species hunt for their food, were obtained from EltonTraits (Wilman et al. 2014), while habitat classifications were obtained from Merchant et al. (2024) and the Animalia database (Animalia 2026). These three datasets were combined into a single habitat/foraging environment category. Fossorial classifications from Merchant et al. (2024) were unchanged, though terrestrial species with fossorial traits reported on the Animalia database (Animalia 2026) were reclassified as ‘semi-fossorial’ to address conflicting information across sources. Arboreal and scansorial strata were merged into an ‘arboreal/scansorial’ category, while ‘marine’, ‘ground’, and ‘aerial’ categories were renamed ‘aquatic’, ‘terrestrial’ and ‘volant’, respectively. Based on taxonomy, all species within Chiroptera (bats) were classified as ‘volant’, Pinnipedia (seals and walruses), Sirenia (manatees and dugongs) and Cetacea (whales and dolphins) as ‘aquatic’, and Ruminantia (ruminant herbivores) as ‘terrestrial’, reflecting the typical ecological habitats of these lineages.

**Figure 1.**
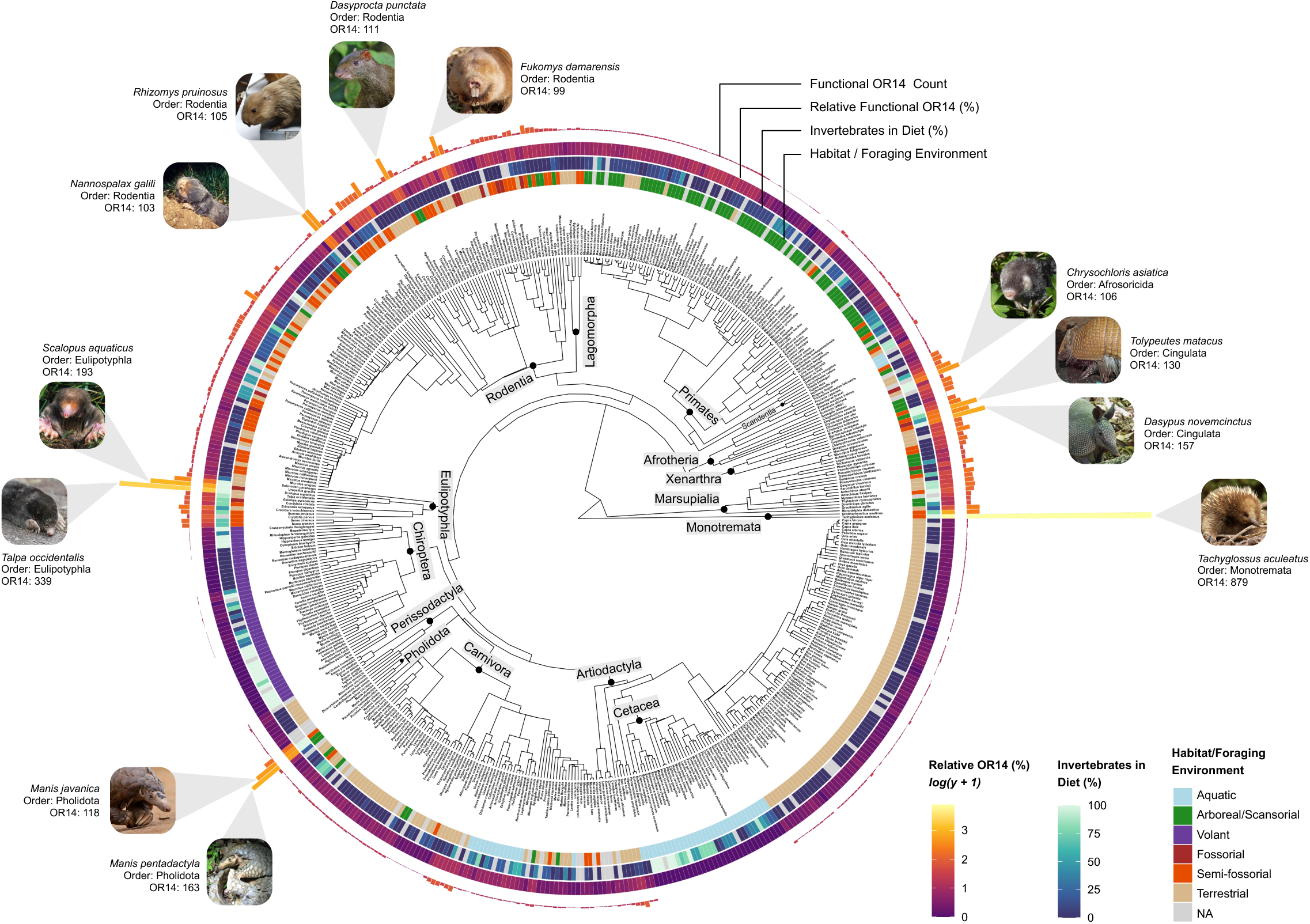
Independent expansion of OR14 across terrestrial and insectivorous mammals. Functional OR14 gene counts are shown as bar plots aligned the mammalian phylogenetic tree. Species with exceptionally high functional OR14 counts (≥99 genes) are highlighted. Relative OR14 proportions are mapped as a heatmap using a continuous colour scale from purple (low) to yellow (high). The percentage of invertebrates in the diet of each species is also mapped, using a gradient from dark blue (low) to light blue (high). Habitat/foraging categories are shown and include aquatic (light blue), arboreal/scansorial (green), volant (purple), fossorial (dark brown), semi-fossorial (orange) and terrestrial (light brown) classifications. Missing data for habitat/dietary traits are shown in grey. Photographs are distributed under a creative commons license (Table S6).

### Investigating the Expansion of OR14 across Terrestrial Insectivores

To test whether functional OR14 counts scale with overall olfactory receptor expansion (Figure 2A) and terrestrial insectivory (Figure 2B), phylogenetic generalized least squares (PGLS) analyses were performed. All PGLS analyses were performed using the caper R package (Orme et al. 2013), with the TimeTree5 (Kumar et al. 2022) phylogenetic tree used to weight residuals. Lambda (λ), which quantifies phylogenetic signal, was estimated for each model using maximum likelihood optimization. A *log(y +1)* transformation was applied to OR14 gene counts to account for skewness, which was assessed using the *e1071* R package (Meyer et al. 2025) (Figure S2). Species were defined as terrestrial insectivores if ≥60% of their diet consists of invertebrates and they inhabit terrestrial environments or exhibit ground-based foraging (Table S2). Species with missing data for either diet or terrestriality categories were excluded from our classifications.

**Figure 2.**
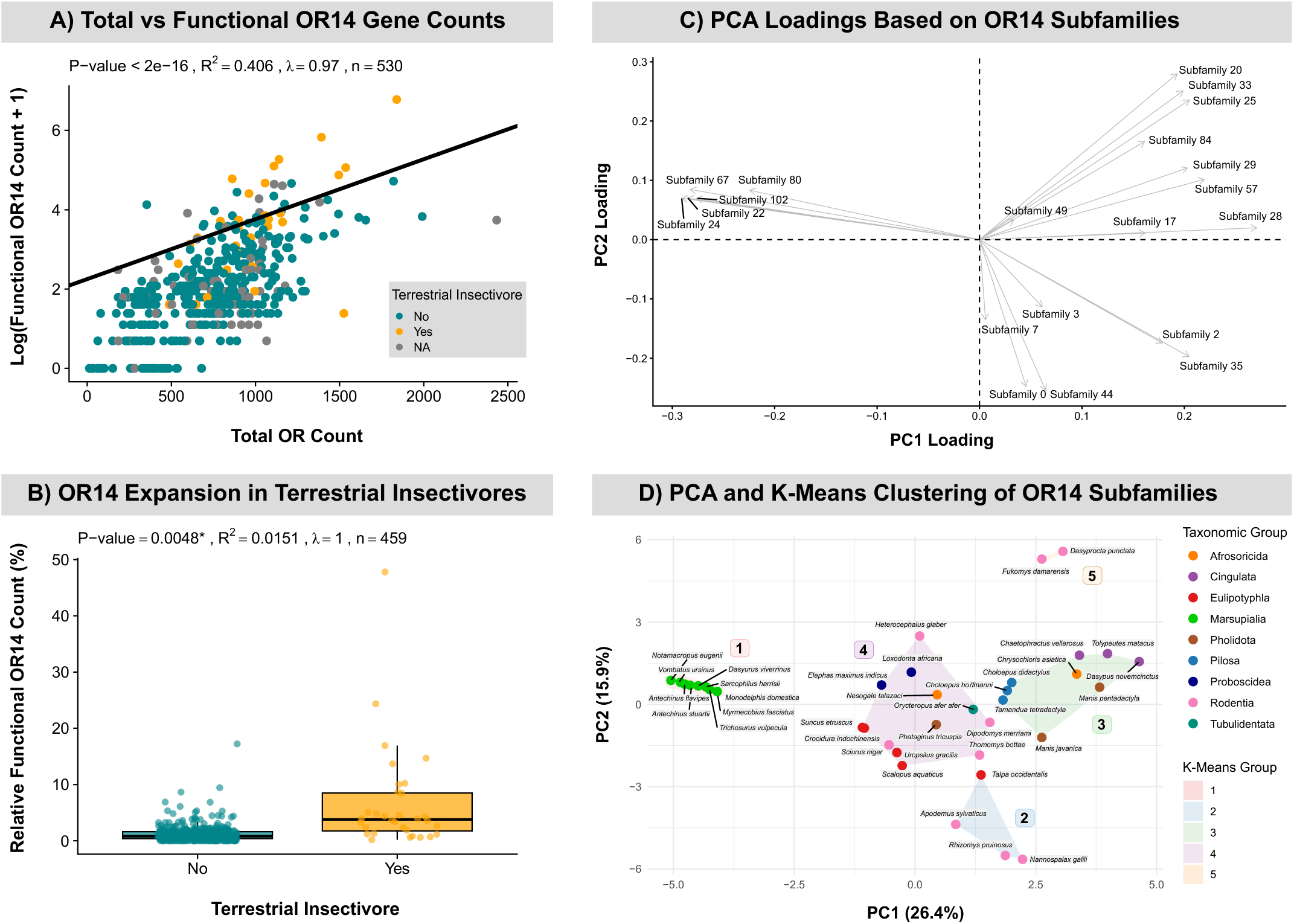
Investigating relative OR14 expansion (A), OR14 expansion in terrestrial insectivores (B), and clustering of species based on OR14 subfamily composition (C, **D)**. Functional OR14 gene counts exhibit a positive and statistically significant relationship with total functional olfactory receptor (OR) repertoire size across mammals (A). After accounting for variation in total OR repertoire size, a significant (p = 0.0048), though weak (R^2^ = 0.0153) association between terrestrial insectivory and relative OR14 repertoire size is observed (B). PCA and K-means clustering reveals divergence across mammals based on OR14 subfamily composition (C, D). PCA loadings are displayed as arrows overlaid on the PCA plotting area, where direction indicates association with principal components and length reflects the magnitude of each subfamily’s contribution (C). PC1 and PC2 together represent just 42.3% of the total variance across all 33 subfamilies (D). K-means cluster groups (Groups 1 -5) were overlaid onto the PCA plot, with species labeled and colored by taxonomic group.

### Principal Component Analysis and K-Means Clustering of OR14 Subfamilies

Principal component analysis (PCA) was performed on OR14 subfamily gene counts to assess whether species occupying similar ecological niches exhibit consistent or divergent patterns of OR14 expansion. OR14 subfamily groups were obtained by clustering functional OR14 sequences at 60% amino acid identity using CD-HIT (Li et al. 2001), with the word size parameter set to 4 (Table S3). An identity value of 60% was chosen based on the conventional minimum threshold for OR subfamily assignment (Glusman et al. 2001, Olender et al. 2020). Only species with ≥40 functional OR14 genes were considered for PCA, to capture species in the upper tail of the OR14 distribution, with a wide range of ecological niches and taxonomic groups represented. OR14 subfamily clusters containing just one species, or with less than 10 representatives across species were excluded from further analysis, as these are more likely to reflect prediction or clustering artefacts, rather than genuine, evolutionarily conserved subfamilies. PCA was performed using the *prcomp* function in R, with OR14 cluster counts adjusted using a *log(y + 1)* transformation. As monotremes are an extreme outlier group in our dataset (Figure S3), PCA was repeated excluding these species to better resolve therian groups (Table S4). PCA loadings were extracted to visualize the contribution of each OR14 subfamily (Figure 2C). To group species based on the PCA results, K-means clustering was applied (Figure 2D), using the *kmeans* function in R, with ‘k’ set to 5 using 25 random starts to ensure stability. The optimal number of clusters (k) was determined using the elbow method based on the total within-cluster sum-of-squares (Figure S4).

## Results

### Expanded OR14 Repertoires in Monotremes and Marsupials

On average, monotreme genomes were found to encode more functional OR14 receptors (mean = 470) than marsupials (mean = 24.52) and eutherians (mean = 11.34). Within Monotremata, *Tachyglossus aculeatus* (Short-beaked echidna) exhibits the largest OR14 repertoire across mammals, with 879 functional receptors (Figure 1, Table S1). While ORs have undergone general expansion in *T. aculeatus,* ranking third among mammals, we find that 47.80% of these functional receptors belong to OR14, indicating a significant bias towards expansion of this family. Similarly, a remarkably high number of OR pseudogenes are reported for this species, of which 63.14% are predicted here to belong to OR14. In comparison, *Ornithorhynchus anatinus* (Platypus) encodes 61 functional OR14 genes (17.23% of total) and 59 OR14 pseudogenes.

Diprotodontia exhibits the widest range of functional OR14 gene counts (Table 1) across marsupials, with a maximum of 65 recorded in *Notamacropus eugenii* (Tammar wallaby) and a minimum of just 14 in *Phascolarctos cinereus* (Koala) and *Pseudochirops corinnae* (Plush-coated ringtail possum). Following *N. eugenii*, the highest functional OR14 counts are observed in *Monodelphis domestica* (Gray short-tailed opossum), *Vombatus ursinus* (Common wombat), and *Dasyurus viverrinus* (Eastern quoll), with each exceeding 49 receptors. Despite having expanded OR14 repertoires, OR14 comprises just 3.02% of total functional ORs in marsupials on average, suggesting that this pattern may reflect general OR expansion rather than enrichment specific to OR14.

**Table 1.** Summary of functional OR14 gene counts across 24 mammalian orders. The mammalian subclass and number of species represented is provided for each order. Summary statistics, including the minimum, maximum, mean, median, and standard deviation of functional OR14 gene counts, are also reported.

| Subclass | Order | No.<br>Species | Min | Max | Mean | Median | Standard<br>Deviation |
| --- | --- | --- | --- | --- | --- | --- | --- |
| Monotremata | Monotremata | 2 | 61 | 879 | 470 | 470 | 578.41 |
| Marsupialia | Dasyuromorphia | 6 | 25 | 49 | 40.33 | 41 | 8.31 |
|  | Didelphimorphia | 2 | 21 | 62 | 41.5 | 41.5 | 28.99 |
|  | Diprotodontia | 12 | 14 | 65 | 31.58 | 30.5 | 16.33 |
|  | Microbiotheria | 1 | 21 | 21 | 21 | 21 | NA |
| Eutheria | Afrosoricida | 3 | 33 | 106 | 70 | 71 | 36.51 |
|  | Artiodactyla | 134 | 0 | 29 | 3.39 | 3 | 3.7 |
|  | Carnivora | 81 | 1 | 26 | 6.44 | 6 | 5.05 |
|  | Chiroptera | 50 | 0 | 3 | 0.52 | 0 | 0.74 |
|  | Cingulata | 3 | 84 | 157 | 123.67 | 130 | 36.91 |
|  | Dermoptera | 2 | 8 | 18 | 13 | 13 | 7.07 |
|  | Eulipotyphla | 12 | 11 | 339 | 74 | 36.5 | 96.58 |
|  | Hyracoidea | 2 | 6 | 7 | 6.5 | 6.5 | 0.71 |
|  | Lagomorpha | 6 | 3 | 11 | 6.67 | 6 | 2.88 |
|  | Macroscelidea | 1 | 17 | 17 | 17 | 17 | NA |
|  | Perissodactyla | 9 | 2 | 9 | 4.22 | 4 | 2.05 |
|  | Pholidota | 4 | 23 | 163 | 88.25 | 83.5 | 63.96 |
|  | Pilosa | 5 | 1 | 57 | 37.8 | 44 | 21.6 |
|  | Primates | 90 | 0 | 15 | 4.7 | 5 | 2.71 |
|  | Proboscidea | 2 | 41 | 45 | 43 | 43 | 2.83 |
|  | Rodentia | 121 | 1 | 111 | 18.98 | 13 | 19.94 |
|  | Scandentia | 3 | 3 | 6 | 4.33 | 4 | 1.53 |
|  | Sirenia | 3 | 3 | 8 | 5.67 | 6 | 2.52 |
|  | Tubulidentata | 1 | 50 | 50 | 50 | 50 | NA |

### Episodic Bursts of OR14 Expansion and Loss Across Eutherians

OR14 has undergone bursts of expansion across eutherian species spanning multiple independent orders (Figure 1). Over 86% of species have OR14 counts lower than the mean, resulting in a right skewed distribution driven by high count outliers (Skewness = 14.37; Figure S2D). Among these outliers, 11 species encoded more than 99 functional OR14 receptors (Figure 1), spanning distantly related orders including Eulipotyphla, Pholidota, Cingulata, Afrosoricida, and Rodentia. The highest eutherian OR14 counts were observed in *Talpa occidentalis* (Iberian mole; 339 receptors) and *Scalopus aquaticus* (Eastern mole; 193), followed by *Manis pentadactyla* (Chinese pangolin; 163) and *Dasypus novemcinctus* (Nine-banded armadillo; 157).

Reduced OR14 repertoires were also observed across several independent clades. Within Chiroptera, 30 of 50 sampled species lacked functional OR14 genes (Table S1). Total loss of the OR14 family was observed in Vespertilionidae, that is, no detectable functional receptors or pseudogenes in the genome of species across this family. Total loss has also occurred in six additional yangochiropteran species, including all sampled species belonging to Miniopteridae (n = 2) and Molossidae (n = 2). A single functional OR14 gene was retained in *E. helvum* and *E. dupreanum*, though no functional OR14 genes were detected in any other yinpterochiropteran species. No pseudogenes were detected in any yinpterochiropteran species except members of the Rousettus genus, each of which contained a single highly degraded pseudogene.

Similarly, no functional OR14 genes were detected across Cetacea. Odontoceti lacked both functional genes and pseudogenes, while a single pseudogene was retained across Mysticeti, except in *Balaenoptera physalus* (Fin whale). *Hexaprotodon liberiensis* (Pygmy hippopotamus) and *Hippopotamus amphibius* (Hippopotamus), the closest extant relatives of cetaceans, both encode 8 functional OR14 genes, placing OR14 loss at the root of the cetacean lineage. Excluding Cetacea, a mean of 6.74 functional receptors was recorded across Artiodactyla, with relative expansion observed within the Suina (pigs) lineage (mean = 17.75).

Within primates, a maximum of 15 functional OR14 receptors was reported in *Daubentonia madagascariensis* (Aye-aye), followed by *Cephalopachus bancanus* (Horsfield’s tarsier) with 14 functional receptors and *Carlito syrichta* (Philippine tarsier) and *Lemur catta* (Ring-tailed lemur) each encoding 11 functional OR14 genes. Diverging patterns are observed within Hominoidea, with a median of 5 functional receptors across Hominidae compared to just 1 receptor in Hylobatidae. A general reduction of OR14 genes has occurred in Platyrrhini, with a median of 1.5 receptors and no functional OR14 genes recorded in *Ateles hybridus* (Brown spider monkey) and *Saimiri boliviensis boliviensis* (Black-capped squirrel monkey).

A mean of 6.44 functional OR14 genes was recorded across carnivorans (Table 1). Relative reduction was observed across Canidae, with each species encoding just 1-2 functional receptors. Unlike fully aquatic cetaceans where no functional receptors were observed, pinnipeds retain 4.17 functional OR14 genes on average. Relative expansion was observed across the Ursidae family, with each species encoding more than 20 functional receptors.

### OR14 Repertoire Expansion in Fossorial and Insectivorous Mammals

Mapping functional OR14 gene counts to diet and habitat revealed a pattern of OR14 expansion among species with convergent adaptations to subterranean or soil-based foraging environments (Figure 1). Among the species with the highest OR14 counts, we observe an overrepresentation of fossorial, semi-fossorial or insectivorous mammals. Notably, these traits span independent orders, and often exhibit convergent morphological features reflective of adaptation to their environments. For example, significant OR14 expansion is observed across true moles (Eulipotyphla), golden moles (Afrosoricida), and multiple mole-rat lineages (Rodentia), many of which exhibit convergent fossorial traits including specialized claws and incisors for digging, and reduced visual anatomy (Figure 1). Similarly, myrmecophagous mammals (species that feed primarily on ants and termites), such as armadillos (Xenarthra), pangolins (Pholidota), and the echidna (Monotremata) exhibit expanded OR14 repertoires and convergent ant-eating adaptations, including long extensible tongues and protective body armor. Further supporting this association, chiropterans exhibit significant reduction or total loss of OR14 repertoires, despite having highly insectivorous diets (Figure 1), likely reflecting their non-terrestrial, volant ecology. Similarly, while aquatic invertebrates make up a significant portion of cetacean and pinniped diets, we observe total loss or reduced OR14 repertoires across these aquatic species. Artiodactyla also exhibit low OR14 counts despite inhabiting terrestrial environments, perhaps consistent with a lack of subterranean foraging or insectivorous diets.

Taken together, these patterns suggest that the combination of terrestrial/fossorial habitats and insectivorous diets is associated with elevated OR14 repertoire sizes. As functional OR14 counts are positively associated with total OR counts (p < 2 x 10^-16^, Figure 2A), we accounted for total repertoire size when testing this association directly. Consistent with our hypothesis, we observe a significant (p = 0.0048), though weak (R^2^ = 0.0151) relationship between terrestrial insectivory and relative OR14 repertoire size (Figure 2B), with a slightly higher mean value in terrestrial insectivores (mean = 2.00%, 18.31 functional OR14) than across other species (mean = 1.32%, 10.43 functional OR14).

### OR14 Subfamilies Segregate with Ecological Niche

Clustering species based on OR14 subfamilies revealed evidence of functional divergence among OR14 repertoires, reflective of adaptation to ecological niche. Including monotremes in our PCA resulted in extreme divergence between monotremes and all other therian species (Figure S3). As such, we find that monotremes exhibit significant sequence divergence across OR14 repertoires relative to marsupials and eutherians. Notably, we also observe subfamily divergence within Monotremata, with clear segregation between *T. aculeatus* and *O. anatinus* recorded. Removing monotremes from a subsequent PCA improved resolution, enabling clear segregation of clusters within Theria to become visible (Figure 2C, D), with the caveat that PC1+PC2 represents just 42.3% of the total variance across all 33 subfamilies (Table S5). We find that marsupial species form a single group based on K-means cluster assignment (Group 1; Figure 2D), divergent from eutherian species. Further, by visualizing the PCA loadings, we report that marsupial species segregate based on unique OR14 subfamilies, specifically, Subfamilies 22, 24, 67, 80 and 102 (Figure 2C).

Within Eutheria, divergence of OR14 repertoires independent of phylogenetic relatedness emerge, indicating that genomic strategies converge among species which occupy similar ecological niches. Highlighting this, myrmecophagous species across Pholidota, Pilosa and Cingulata cluster together (Group 3, Figure 2D), despite representing divergent taxonomic orders. *Chrysochloris asiatica* (Cape golden mole) also lies within this group, segregating from *Nesogale talazaci* (Talazac’s shrew tenrec) despite both belonging to Afrosoricida. OR14 subfamilies 17, 28, 29 and 57 primarily drive the separation of this group in the PCA (Figure 2C). Interestingly, *T. aculeatus* and *Orycteropus afer afer* (Southern aardvark) do not cluster in this group, despite being myrmecophagous. Conversely, it is unclear why sloths (*Choloepus hoffmanni* and *C. didactylus*) group within this cluster, though this may reflect phylogenetic relatedness across Xenarthra.

While most members of Eulipotyphla group together (Group 4, Figure 2D), *T. occidentalis* forms a distinct cluster with three rodent species, namely *Apodemus sylvaticus* (Wood mouse), *Rhizomys pruinosus* (Hoary bamboo rat) and *Nannospalax galili* (Middle East blind mole-rat) (Group 3, Figure 2D). Subfamilies 0, 3, 7 and 44 drive separation of this group (Figure 2C). Rodent species are also distributed across K-means Group 4 and 5, with Group 5 forming a rodent-specific cluster and showing strong separation from other groups based on subfamilies 20, 25 and 33. Specifically, *Dasyprocta punctata* (Central American agouti) and *Fukomys damarensis* (Damara Mole-rat) are represented in this group, belonging to the Dasyproctidae and Bathyergidae families. Notably, these species are separated from *Heterocephalus glaber* (Naked mole-rat; Group 4), also a member of the Bathyergidae family, indicating divergent OR14 repertoires within the Hystricomorpha suborder. Other rodent species which lie in Group 4 include *Sciurus niger* (Fox squirrel), *Dipodomys merriami* (Merriam’s kangaroo rat) and *Thomomys bottae* (Botta’s pocket gopher), representing the Sciuromorpha and Castorimorpha suborders. Group 4 also includes species across Tubulidentata, Proboscidea, Afrosoricida, Pholidota, and Eulipotyphla, forming a heterogeneous cluster with a broad range of phenotypically and ecologically diverse lineages.

## Discussion

This study represents the first large-scale investigation into the evolution of the OR14 gene family across hundreds of mammalian species. By including a taxonomically diverse dataset, representing a broad range of diets, habitats, and life-history traits, we uncover new insights into the ecological factors driving OR14 variation. Episodic bursts of OR14 expansion were observed across multiple independent lineages, with fossorial and insectivorous mammals disproportionately represented. Additionally, OR14 subfamilies showed evidence of divergence among species independent of phylogeny, consistent with niche specialization. Together, our results suggest that OR14 expansion may arise through ecological convergence, reflecting olfactory adaptations associated with insectivorous and fossorial lifestyles.

### Evolution of OR14 Genes Across Mammals

The number of OR14 genes varied across mammalian lineages, with eutherians typically having fewer functional OR14 genes than monotremes and marsupials. The generally expanded OR14 repertoires across marsupials and monotremes may be indicative of their importance in the ancestral mammalian state (Sampson et al. 2024). Notably, the last common ancestor of mammals is widely hypothesized to have been insectivorous (Brocklehurst et al. 2022, Gerkema et al. 2013, Wu et al. 2021), raising the possibility that high OR14 copy numbers in these lineages may indicate some degree of conservation of an ancestral repertoire, particularly as many species across these clades are also insectivorous. The contrasting gene family size across most placental mammals may represent a shift towards alternative food sources. This hypothesis is supported by convergent episodic bursts of OR14 gene expansion across several terrestrial insectivorous mammalian lineages, where copy numbers can reach up to ∼30 times the eutherian mean. The repeated expansion of OR14 repertoires across these independent lineages suggests that OR14 duplication and retention is adaptive, in the same way that OR5/8/9 has been linked to herbivory in mammals (Hughes et al. 2018) or OR1/3/7 to frugivory in bats (Hayden et al. 2014).

### Convergent Signals of Insectivory and Fossorial Adaptation

The independent expansion of OR14 across myrmecophagous and mole-like mammals is notable, as these groups also exhibit convergence at both morphological and ecological levels (Cheng et al. 2023, Partha et al. 2017). As such, expansion of this gene family may also be convergent, reflecting some functional role in insectivory and/or subterranean habitats. This pattern is further supported by the total loss of OR14 genes across cetaceans and multiple bat lineages, potentially arising under relaxed selective pressure coinciding with transitions away from ground or soil-based foraging. Consistent with this, we find a statistically significant association between relative OR14 gene counts and terrestrial insectivory. However, we note that the explained variance is low, possibly resulting from our definition being too broad, or due to outlier species that cannot be explained by this model. For example, *Chaetophractus vellerosus* (Screaming hairy armadillo) is fossorial (Poljak et al. 2018), exhibits high OR14 counts, and is insectivorous, but falls just below the 60% invertebrate diet threshold used here to define terrestrial insectivores (Wilman et al. 2014). Similarly, while *Tamandua tetradactyla* (Southern tamandua) is a primary insectivore (Deloss et al. 2024), its arboreal/scansorial foraging pattern (Wilman et al. 2014) excludes it from our definition. We also observe exceptions in species such as *Dasyprocta punctata* (Central American agouti) and *Loxodonta africana* (African elephant) which exhibit high OR14 gene counts despite neither species being fossorial nor primarily insectivorous.

### OR14 Subfamily Convergence in Myrmecophagous Mammals

Given the lack of visual or auditory environmental cues in subterranean environments, a well-developed sense of olfaction is crucial in fossorial species (Catania 2005, Heth et al. 2002, Ruiz-Rubio et al. 2024). Similarly, olfaction is an important method for detecting insects such as ants and termites in nests/tunnels that are otherwise visually concealed (DiPaola et al. 2020, Ocko et al. 2019). It is unsurprising therefore that we observe expanded olfactory repertoires across these species. While both myrmecophagous and mole-like fossorial species converge on OR14 expansion, perhaps unified by shared soil-based foraging environments, their diets can vary substantially in composition (Bolković et al. 1995, Carlini et al. 2016, Fielden et al. 2009, García-López de Hierro et al. 2013, Hartman et al. 2000, Sikes et al. 1990, Tamang et al. 2022, Zhang et al. 2019). We therefore expected to see some level of sequence divergence among receptor subfamilies across these groups. Supporting our hypothesis, we find evidence of genomic convergence across many phylogenetically independent myrmecophagous mammals (except *T. aculeatus* and *O. afer afer*), with expansion of similar OR14 subfamilies grouping these lineages together. Specific OR14 subfamilies may be under selection in parallel, with receptors tuned specifically to detect odorants associated with ants and termites. Interestingly, *Chrysochloris asiatica* (Golden mole) also lies in this group, segregating from other mole-like species. While there is little information regarding the dietary composition of *C. asiatica* directly, beyond general insectivory (Bennett and Spinks 1995), termites are known to make up the majority of the diet of *Eremitalpa granti namibensis* (Grant’s golden mole), a closely related member of the Chrysochloridae family (Fielden et al. 1990).

### Functional Divergence of OR14 Genes

Monotremes and marsupials showed extreme divergence from eutherian species based on OR14 subfamily composition. Within monotremes, divergence is also observed at the species level, segregating *O. anatinus* from *T. aculeatus*, which has unusually extensive duplication and pseudogenization rates. These high duplication rates may be indicative of explorative expansion of the OR14 gene family associated with species-specific adaptations. Despite this, *T. aculeatus* does not cluster with other myrmecophagous mammals (Sprent and Nicol 2016). It is possible that high OR14 divergence in marsupials and monotremes reflects neofunctionalization, with ectopic expression of OR14 receptors in T-cells reported in these lineages suggesting diversification of function towards a role in immunity (Sampson et al. 2024). There is also indirect evidence that OR14 may play a role in immunity in humans, where variation in OR14J1 has been associated with type 1 diabetes (Jahromi 2016, Jahromi et al. 2014). Despite encoding only seven functional OR14 genes, the human OR14 repertoire has the lowest mutation rate in comparison to other OR families (Jimenez et al. 2021), consistent with some form of functional constraint. It remains unclear as to why OR14 genes with putative immune-related functions would exhibit independent expansion across insectivorous and subterranean mammals, outside of their role in olfaction.

## Conclusion

The sense of olfaction, encoded by the OR gene repertoire, provides a key means of accessing sensory information related to diet and habitat in mammals. Our comprehensive analysis of the OR14 gene family across 555 species reveals an association between ecological niche adaptation and sensory gene expansion. While OR14 is typically reduced in eutherians compared to monotremes and marsupials, we show that it has undergone repeated expansion in multiple independent lineages, driven in part by both insectivorous diets and fossorial habitats. Our findings provide insight into the role of sensory perception in mammalian evolution, highlighting the importance of specific gene subfamilies in adaptation to diverse ecological niches.

## Data availability

Olfactory receptor gene predictions generated in this study are available via Zenodo at the following link: (https://zenodo.org/records/21261236?preview=1&token=eyJhbGciOiJIUzUxMiJ9.eyJpZCI6ImNiYTcyNTgxLTA1MjUtNDJkZC1hYTAwLWYzMTY3MmU3MzNkZCIsImRhdGEiOnt9LCJyYW5kb20iOiIwZmRhY2NmYWNmMjE1OGY4NzU4NjliNzIwMjdjMmI2MCJ9.9e-uthzDUUfILChVvGLOg2tyI-sh04AoquzFyFtys7PUD2QuAFvHnvPM_pPvhh-m5QffHTkViWJzmLlpafRAlA).

## Acknowledgments

Images of all species included in this publication are distributed under the creative commons license and were accredited where available (Table S6).

## Author Contributions

Conceptualization of the study, L.R. and G.M.H. Methodology and study design, L.R. and G.M.H. Sensory receptor gene mining and classification, L.R. Data analysis, interpretation, and visualization, L.R. Writing, reviewing, and editing the manuscript, L.R. and G.M.H. Supervision and project oversight, G.M.H. Both authors have read and approved the final manuscript.

## Supplementary material

Supplementary tables and figures are available online.

## Supplementary Figures

Figure S1. Original uncorrected and manually corrected OR14 prediction from *Eidolon helvum,* using the orthologous sequence from sister taxa *Eidolon dupreanum* to guide annotation. The original prediction (red) starts at a downstream, in-frame, methionine residue, resulting in a truncated prediction relative to the orthologous *E. dupreanum* sequence. Correcting this prediction manually using TBLASTN results in an updated classification, where both sister taxa encode a full length OR14 protein (blue).

**Figure S2. Distribution of functional OR gene counts before and after *log(y +1)* transformation.** Transforming total functional OR counts increases skewness (A, B). Therefore, total functional OR counts were not transformed in this study. In contrast, functional OR14 gene counts were highly positively skewed prior to transformation, with skewness decreasing from 14.37 (C) to 0.41 following *log(y + 1)* transformation (D). Functional OR14 counts were therefore transformed for all subsequent analyses to improve normality.

**Figure S3. Principal Component Analysis (PCA) based on OR14 subfamily composition, including monotremes.** Only species with ≥40 functional OR14 genes were considered for PCA. Including monotremes illustrates extreme divergence between this clade and all other therian species, making it difficult to resolve clusters across therian species. Significant divergence is observed within monotremes, with clear segregation between Tachyglossus aculeatus and Ornithorhynchus anatinus recorded.

**Figure S4. Elbow plot used to determine the optimal number of clusters (*k*) for k-means clustering.** Using the elbow method, based on the total within-cluster sum of squares, the number of clusters (*k*) was set to 5.

## Supplementary Tables

**Table S1. Summary of OR gene counts across 555 mammalian species used in this study.** Genome accession IDs are provided for each species, along with phylogenetic classifications, and OR gene counts.

**Table S2. Habitat and dietary classifications for each species used in this study.** Habitat classifications from each data source are provided, along with the final merged habitat/foraging environment classification used in this study. The percentage of invertebrates in the diet of each species is also provided, along with the terrestrial insectivore classifications assigned in this study.

**Table S3. OR14 subfamily counts across all species.** Subfamily classifications were derived using CD-HIT with a 60% sequence identity threshold. CD-HIT cluster identifiers were retained and used to define subfamilies (i.e Cluster 1, was renamed as Subfamily 1).

**Table S4. OR14 subfamilies included in the principal component analysis. A**fter filtering to include only species with ≥40 functional OR14 genes and OR14 subfamilies represented by more than one species and with more than 10 functional genes across all species, a total of 33 subfamilies remained for analysis.

**Table S5. Principal component analysis summary.** The standard deviation, proportion of variance explained, and cumulative proportion for each principal component (PC) are provided.

**Table S6. Creative Commons licensing and author attribution for each photograph used in this study**. A link to the original image and its license is provided, along with details of any image modifications.

