## Supplementary Figures for "Expansion of Olfactory Receptor Family 14 Across Fossorial and Insectivorous Mammals"

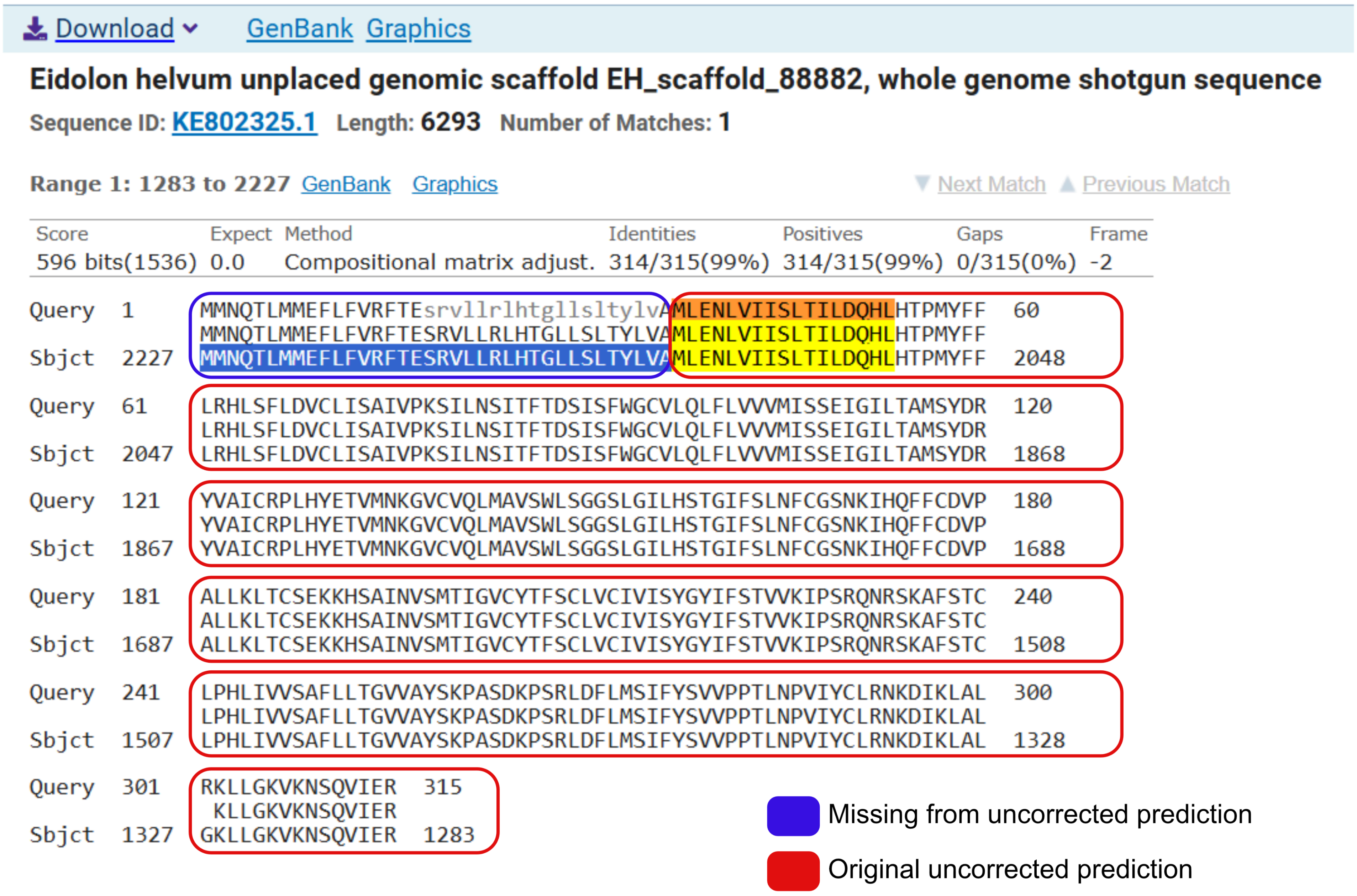


**Figure S1. Original uncorrected and manually corrected OR14 prediction from *Eidolon helvum*, using the orthologous sequence from sister taxa *Eidolon dupreanum* to guide annotation.** The original prediction (red) starts at a downstream, in-frame, methionine residue, resulting in a truncated prediction relative to the orthologous *E. dupreanum* sequence. Correcting this prediction manually using TBLASTN results in an updated classification, where both sister taxa encode a full length OR14 protein (blue).

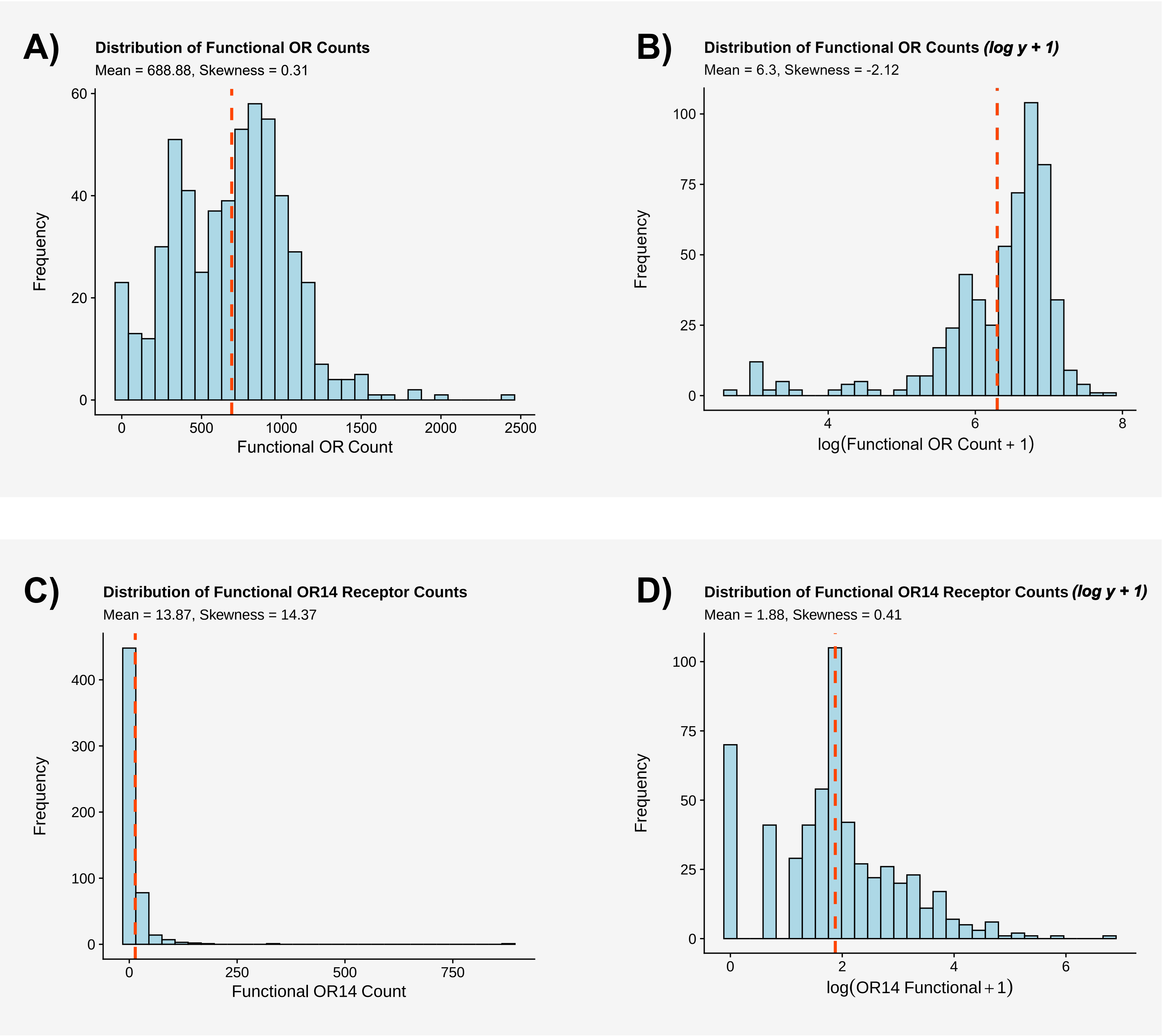


**Figure S2. Distribution of functional OR gene counts before and after *log(y + 1)* transformation.** Transforming total functional OR counts increases skewness (A, B). Therefore, total functional OR counts were not transformed in this study. In contrast, functional OR14 gene counts were highly positively skewed prior to transformation, with skewness decreasing from 14.37 (C) to 0.41 following *log(y + 1)* transformation (D). Functional OR14 counts were therefore transformed for all subsequent analyses to improve normality.

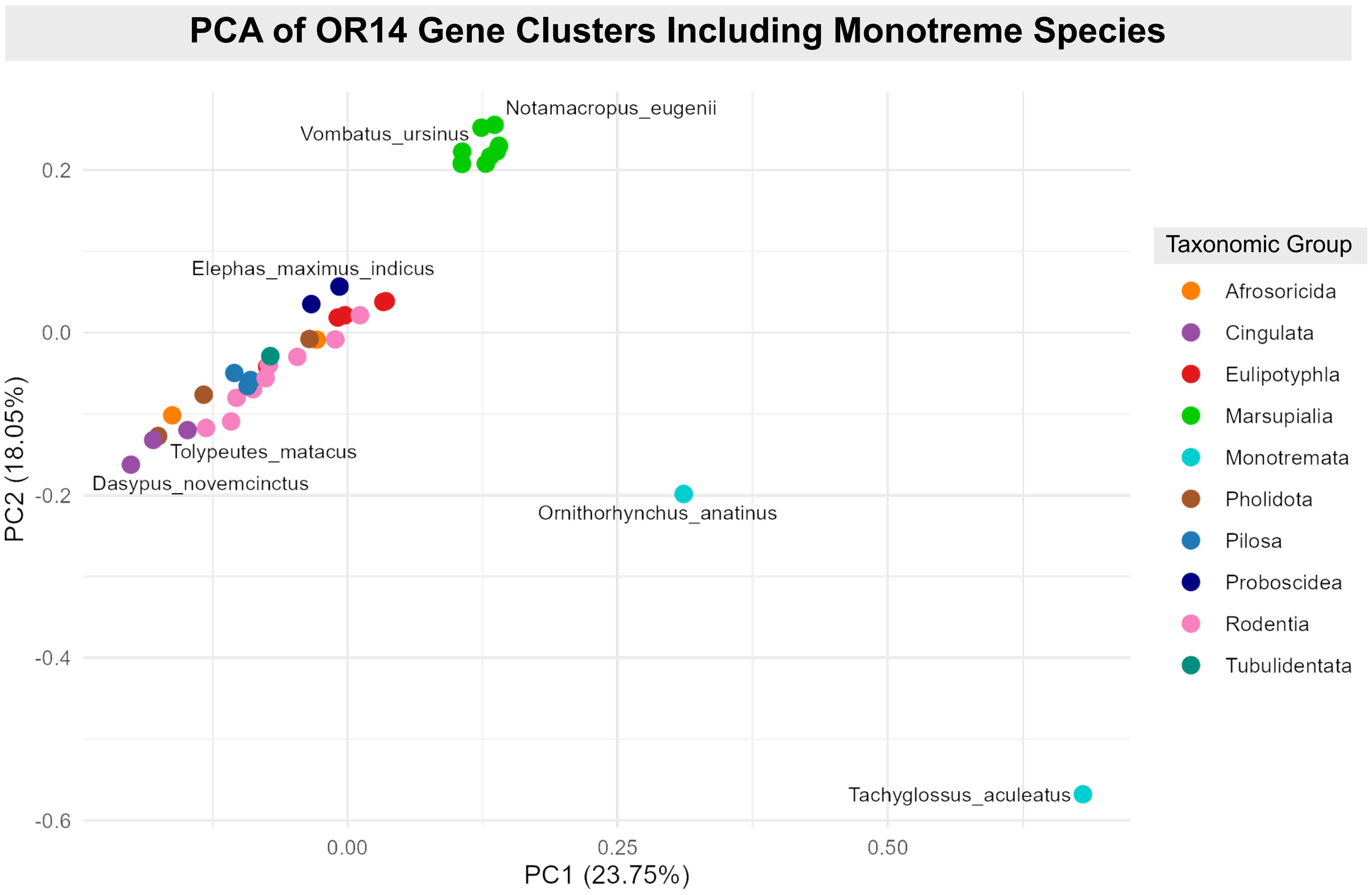


**Figure S3. Principal Component Analysis (PCA) based on OR14 subfamily composition, including monotremes.** Only species with ≥40 functional OR14 genes were considered for PCA. Including monotremes illustrates extreme divergence between this clade and all other therian species, making it difficult to resolve clusters across therian species. Significant divergence is observed within monotremes, with clear segregation between *Tachyglossus aculeatus* and *Ornithorhynchus anatinus* recorded.

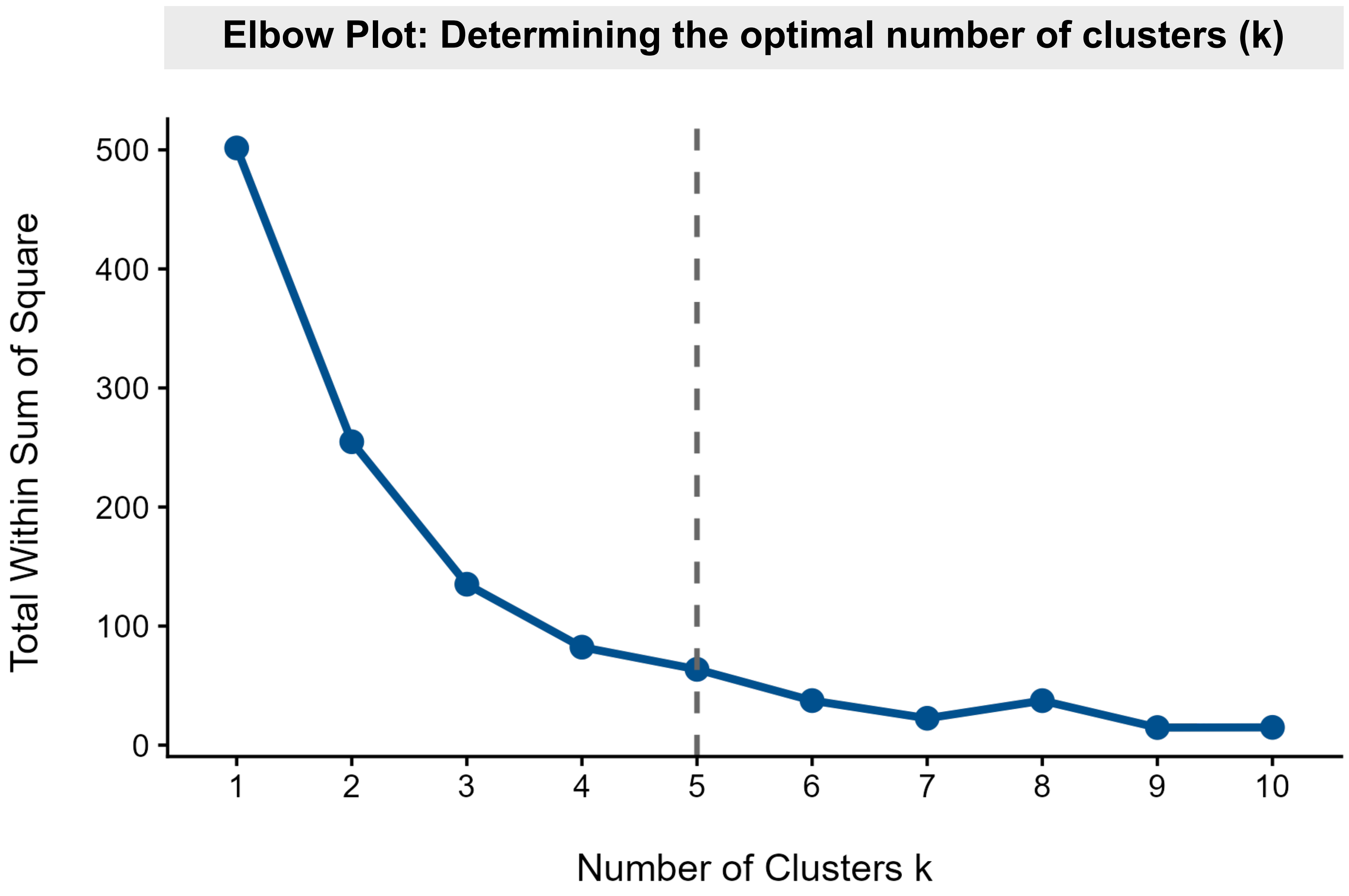


**Figure S4. Elbow plot used to determine the optimal number of clusters (k) for k-means clustering.** Using the elbow method, based on the total within-cluster sum of squares, the number of clusters (k) was set to 5.
